# Elimination of the neuroparsin neuroendocrine cells in *Drosophila virilis* using the UAS-Gal4 system shows that neuroparsin is not important for reproduction in this species

**DOI:** 10.1101/2024.08.30.610547

**Authors:** Jan A. Veenstra

## Abstract

Neuroparsin is a common insect neurohormone produced in large neuroendocrine cells in the brain and is important in mosquito reproduction. Although it is present in many flies including many *Drosophila species*, it was lost from *D. melanogaster* and a few closely related species. Three different lines of transgenic *D. virilis* were produced: One that expresses the yeast transcription factor gal4 under the control of the neuroparsin promoter (NP-gal4), while others codes for proteins under the control of the gal4 promoter, either enhanced green fluorescent protein (UAS-eGFP) or the *D. virilis* ortholog of reaper, an apoptosis inducing protein (UAS-rpr). Crosses between UAS-eGFP and NP-gal4 revealed that expression of NP-gal4 was correct. Crosses between UAS-rpr and NP-gal4 completely eliminated the neuroparsin neuroendocrine cells, but were without effect on reproduction.

## Introduction

Neuroparsin is a hormone that is commonly present in insects and decapods (e.g. Veenstra, 2010, 2016). Several species have a number of paralog neuroparsin genes, while in migratory locusts a single gene produces five different neuroparsin transcripts, each coding for a slightly different neuroparsin (Veenstra, 2010, 2016). Unlike most neuropeptides and neurohomones it does not act through a G-protein coupled receptor but it stimulates a venus kinase receptor that has a tyrosine kinase domain similar to the one in insulin and IGF receptors (Vogel et al., 2015). Evidence in locusts suggests that neuroparsin antagonizes the action of juvenile hormone (Girardie et al., 1987), which is important for vitellogenesis. Indeed, inhibiting neuroparsin expression by RNAi increases expression of vitellogenin and slightly accelerates oocyte growth in this species (Badisco et al., 2011). In ant and bumblebee workers vitellogenin production and neuroparsin are negatively correlated and more recently it was shown that in ants neuroparsin increases the production of juvenile hormone binding protein and decreases that of shadow, one of the enzymes in the cascade of the juvenile hormone synthesis (Nagel et al., 2020; Santos et al., 2022; Zhang et al., 2024). In at least some decapods neuroparsin similarly inhibits vitellogenin expression (Yang et al., 2014; Liu et al., 2020). Nevertheless, in mosquitoes neuroparsin is responsible for initiating vitellogenesis through the stimulation of the synthesis of ecdysone by the ovary (Brown et al., 1998). Thus it is difficult to discern a unified function for this ubiquitous hormone. Surprisingly, whereas it is present in many *Drosophila* species, it is absent from *D. melanogaster* (Veenstra, 2010), the insect model *par excellence*.

Piggybac transposable elements can be used to introduce transgenes in insects and the Gal4-UAS tools that are extensively used in *D. melanogaster* to fine tune transgene expression, have been reported to be effective also in other *Drosophila* species (Handler, 2002; Holtzman et al., 2010). The apparent contradiction between the absence of neuroparsin in *D. melanogaster* and its essential function in mosquitoes suggested that it would be interesting to know more about this hormone in another *Drosophila* species, one that is not too closely related to *D. melanogaster*, like *D. virilis*.

## Materials and Methods

### Antisera

Antisera were raised to FTIPNYED, the C-terminus of *D. virilis* neuroparsin, and GFNRLIPQ, the C-terminal sequence of the B-chain of *D. virilis* Ilp2. Peptides were custom synthetized by Genecust (Boynes, France) and coupled to bovine serum albumin using 1,5-difluoro-2,4-dinitrobenzene as described by Tager (1976). These conjugates were used to raise antisera in both mice and rabbits (Yorkshire Bioscience Ltd,York, UK). Rabbit antiserum to neuroparsin and mouse antiserum to ilp2 were used here.

### Transgenic *D. virilis* flies

Piggybac vector pXLBacII/PubRFP.T3 was a generous gift from Al Handler (USDA, Gainesville, Florida 32608, USA). This vector uses a fluorescent marker which at second thought was considered undesirable and replaced with the miniwhite gene from pUAST. Piggybac-Gal4 and Piggybac-UAST vectors were then constructed by adding the corresponding sequences from pUAST and pGaTB. The *D. virilis* neuroparsin promoter was amplified from *D. virilis* genomic DNA using 5’-GGATGGATCCTCTGGGAAGGTACGTGGAAC-3’ and 5’-GGTTGCGGCCGCTAGGATAACTGCTTCTGCTGGGCAACAT-3’ and cloned into the Piggybac-Gal4 vector. The eGFP coding sequence was amplified from *D. melanogaster* UAS-eGFP flies using 5’-GGACCGAATTCAGGCCTGTTTAAACGATCCAC-3’ and 5’-GGACCGAGATCTGGAGGTGTGGGAGGTTTTT-3’ and cloned into the Piggybac-UAST vector, while 5’-GGGAGAATTCCAACGCCAACAACAAGAAGA-3’ and 5’-GGAAGCGGCCGCTTTTTCACTGCGATGGTTTG-3’ were used to amplify the *D. virilis* ortholog of the *D. melanogaster* rpr gene and also cloned into the Piggybac-UAST vector. Since the Piggybac-UAS-rpr was unsatisfactory, a second Piggybac-UAS-rpr was produced using the translation promoters and the 10XUAS sequence that are present in the pJFRC29-10XUAS-IVSmyr::GFP-p19 transgene (Pfeiffer et al., 2012). The correct sequences of the various constructs were confirmed by Sanger sequencing. Fly transformation was mostly done by Rainbow Transgenic Flies Inc. (Camarillo, CA 93012 USA), but the neuroparsin-gal4 lines were make by Genetic Services Inc. (Sudbury, MA 01776 USA).

### Flies

Flies were maintained on standard fly food, *i*.*e*. yeast extract and corn meal with agar. *D. virilis* lacks balancers, making flies homozygous thus requires being able to distinguish between hemi- and homozygotes which was achieved using eye color.

## Results and Discussion

After injection of *D*.*virilis* embryos a single transgenic fly was obtained that contained two insertions on the XXX chromosome. These two insertions separated immediately and may thus have been on different chromosomes. A much larger number of independent insertions of UAS-eGFP and UAS-rpr were obtained. Although the crosses between the neuroparsin-gal4 and all UAS-eGFP always labeled the neuroparsin neuroendocrine cells, crosses between the neuroparsin-gal4 and UAS-rpr strains, with one exception, did not eliminate the neuroparsin neuroendocrine cells. The exception was a cross in which a transgenic male that was crossed with virgin neuroparsin-gal4 flies, produced some progeny in which the neuroparsin cells were lacking. This male later turned out to have several UAS-rpr insertions in its genome. This suggested that either the *D. virilis* reaper ortholog is less potent than *D. melanogaster* reaper, or that the translation of the UAS-rpr transgene is too weak. The latter hypothesis is supported by the relative weak expression of the UAS-eGFP transgene (as compared to eGFP expression in *D. melanogaster*). I thus tried to increase translation of the UAS-rpr transgene by increasing the number of UAS sequences from five to ten and adding translational enhancers that were shown to be effective in *D. melanogaster* (Pfeiffer et al., 2010). This new UAS-transgene effectively eliminates the neuroparsin neuroendocrine cells.

**Fig 1.**
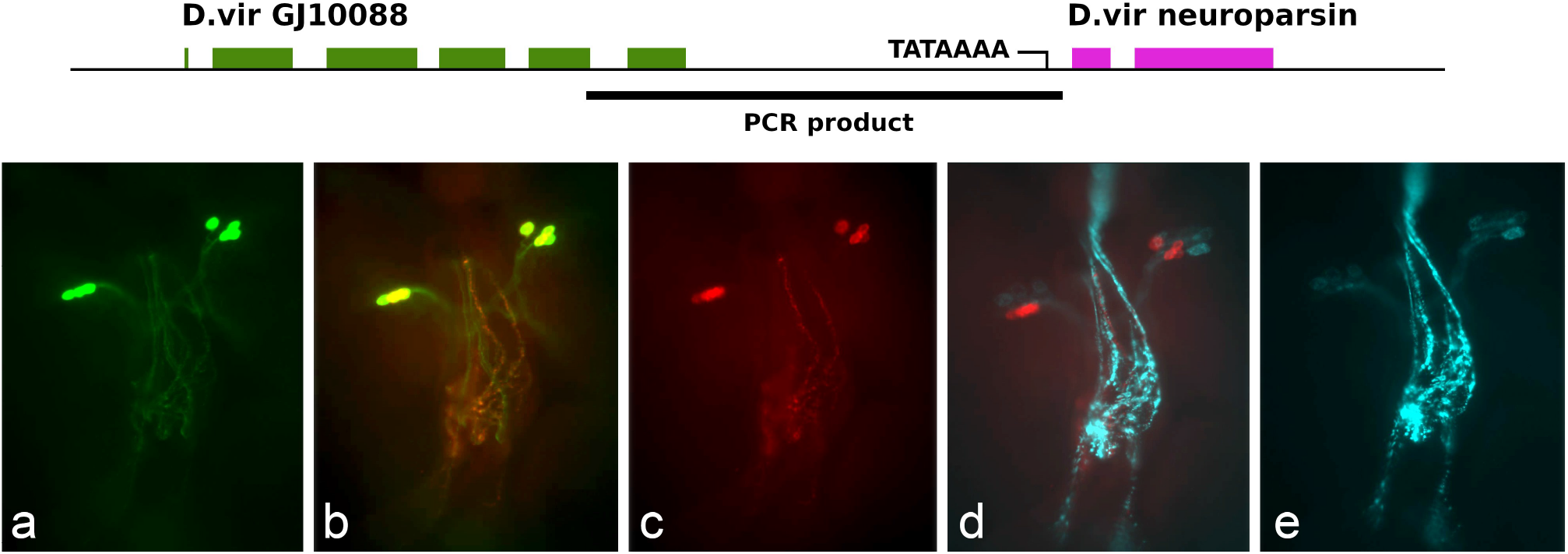
Top: Location of the DNA fragment used to drive expression of gal4 in the neuroparsin-gal4 gene. The tick underline indicates the PCR amplified region. At the 5’-end it encompasses the last coding exon of the GJ10088 gene and at the 3’-end it includes the TATA box and the transcription start site of the neuroparsin gene. Bottom: Expression of eGFP in four bilateral neuroendocrine cells in a neuroparsin-gal4/UAS-eGFP larva (green in a and b). These are the same cells as those labeled by *D. virilis* neuroparsin antiserum (red in b and c). The neuroparsin cells are located next to those producing insulin-like peptides, visualized with an antiserum to vilp2 (cyan in d and e). Note that the neuroparsin immunoreactive projections in the corpora cardiaca are not as prominent as the vilp2 immunoreactive ones. Immunoreactivity of the vilp2 cell bodies is relatively weak as the epitope selected for antibody production only becomes accessible after proteolytic processing of the vilp2 precursor, which is a relatively slow process occurring during transport of the secretory granules from the cell bodies to their release sites in the *corpora cardiaca*.

The expression of the two independent neuroparsin-gal4 lines were identical. They express gal4 in four bilateral brain neuroendocrine cells in both larvae and adults as shown in progeny when these flies are crossed with one of each of ten UAS-eGFP that were kept (Fig. 2a). Rabbit antiserum raised to the C-terminal of *D. virilis* neuroparsin labels the same cells (Figs. 2b, 2c). The neuroparsin cells are located next to seven neuroendocrine cells that are immunoreactive with antiserum to *D. virilis* ilp2 (Figs. 2d, 2e) and like the latter project to the *corpora cardiaca*. The neuroparsin cells are smaller in size and number as the cells labeled by the ilp2 antiserum, while their axon projections into the retrocerebral neuroendocrine complex are much reduced as compared to those of the dilp2 cells.

The reproduction of flies carrying both a neuroparsin-gal4 and a UAS-rpr transgene - and thus lacked the neuroparsin neuroendocrine cells -appeared normal. Although no extensive experiments were performed, in preliminary assays eggs, larvae and pupae all appeared at the same time and in the same quantities as in controls.

Flies lacking neuroparsin neuroendocrine cells reproduced normally, suggesting that under laboratory conditions the neuroparsin cells are not essential. This is reminiscent of the silk worm *Bombyx mori*, where neuroparsin has evolved into a pseudogene in many of the strains that are exploited for the production of silk. Other domesticated *B. mori* strains as well as its close relative *B. mandarina* seem to have a functional neuroparsin gene, like all other Lepidoptera that have been studied (Veenstra, 2020). This might suggest that some insects no longer need neuroparsin when cared for and protected by humans. It is interesting in this respect that in *Anopheles coluzzi* the neuroparsin receptor is important for protection against infection by *Plasmodium* parasites (Gouignard et al., 2019).

This work was financed by institutional funding from the CNRS.

## References

Badisco, L., Marchal, E., Van Wielendaele, P., Verlinden, H., Vleugels, R., Vanden Broeck, J. 2011. RNA interference of insulin-related peptide and neuroparsins affects vitellogenesis in the desert locust Schistocerca gregaria. Peptides 32, 573–80.

Brown, M.R., Graf, R., Swiderek, K.M., Fendley, D., Stracker, T.H., Champagne, D.E., Lea, A.O. 1998. Identification of a steroidogenic neurohormone in female mosquitoes. J. Biol. Chem. 273, 3967–71.

Girardie, J., Boureme, D., Couillaud, F., Tamarelle, M., Girardie, A. 1987. Anti-juvenile effect of neuroparsin A, a neuroprotein isolated from locust corpora cardiaca. Insect Biochemistry 17, 977–983.

Gouignard, N., Cherrier, F., Brito-Fravallo, E., Pain, A., Zmarlak, N.M., Cailliau, K., Genève, C., Vernick, K.D., Dissous, C., Mitri, C. 2019. Dual role of the Anopheles coluzzii Venus Kinase Receptor in both larval growth and immunity. Sci. Rep. 9, 3615.

Grönke, S., Clarke, D.F., Broughton, S., Andrews, T.D., Partridge, L. 2010. Molecular evolution and functional characterization of Drosophila insulin-like peptides. PLoS Genet. 6, e1000857.

Handler, A.M. 2002. Use of the piggyBac transposon for germ-line transformation of insects. Insect Biochem Mol Biol 32, 1211–20.

Holtzman, S., Miller, D., Eisman, R., Kuwayama, H., Niimi, T., Kaufman, T. 2010. Transgenic tools for members of the genus Drosophila with sequenced genomes. Fly (Austin) 4, 349–62.

Liu, J, Liu, A., Liu, F., Huang, H., Ye, H. 2020. Role of neuroparsin 1 in vitellogenesis in the mud crab, Scylla paramamosain. Gen. Comp. Endocrinol. 285, 113248.

McGuire, S.E., Roman, G., Davis, R.L. 2004. Gene expression systems in Drosophila: a synthesis of time and space. Trends Genet. 20, 384–91.

Nagel, M., Qiu, B., Brandenborg, L.E., Larsen, R.S., Ning, D., Boomsma, J.J., Zhang, G. 2020. The gene expression network regulating queen brain remodeling after insemination and its parallel use in ants with reproductive workers. Sci. Adv. 6, 5772.

Pfeiffer, B.D., Truman, J.W., Rubin, G.M. 2012. Using translational enhancers to increase transgene expression in Drosophila. Proc. Natl. Acad. Sci. USA 109, 6626–31.

Santos, P.K.F., Galbraith, D.A., Starkey, J., Amsalem, E. 2022. The effect of the brood and the queen on early gene expression in bumble bee workers’ brains. Sci. Rep. 12, 2018.

Tager, H.S. 1976. Coupling of peptides to albumin with difluorodinitrobenzene. Anal. Biochem. 71, 367–75.

Veenstra, J.A. 2010. What the loss of the hormone neuroparsin in the melanogaster subgroup of Drosophila can tell us about its function. Insect Biochem. Mol. Biol. 40, 354–61.

Veenstra, JA. 2014. The contribution of the genomes of a termite and a locust to our understanding of insect neuropeptides and neurohormones. Front. Physiol. 5, 454.

Veenstra, J.A. 2016. Similarities between decapod and insect neuropeptidomes. PeerJ 4, e2043.

Veenstra, J.A. 2020. Most lepidopteran neuroparsin genes seem functional, but in some domesticated silkworm strains it has a fatal mutation. Gen. Comp. Endocrinol. 285, 113274.

Vogel, K.J., Brown, M.R., Strand, M.R. 2015. Ovary ecdysteroidogenic hormone requires a receptor tyrosine kinase to activate egg formation in the mosquito Aedes aegypti. Proc. Natl. Acad. Sci USA 112, 5057–62.

Yang, S.P., He, J.-G., Sun, C.B., Chan, S.F. 2014. Characterization of the shrimp neuroparsin (MeNPLP): RNAi silencing resulted in inhibition of vitellogenesis. FEBS Open Bio 4, 976–86.

Zhang, X., Xie, N., Ding, G., Ning, D., Dai, W., Xiong, Z., Zhong, W., Zuo, D., Zhao, J., Zhang, P., Liu, C., Li, Q., Ran, H., Liu, W., Zhang, G., 2024. An evolutionarily conserved pathway mediated by neuroparsin-A regulates reproductive plasticity in ants. PLoS Biol. 22, e3002763. 10.1371.

